# Red Light Mediated Photo-Conversion of Silicon Rhodamines to Oxygen Rhodamines for Single-Molecule Microscopy

**DOI:** 10.1101/2024.05.21.595223

**Authors:** Jacob M. Ritz, Aset Khakimzhan, Joseph J. Dalluge, Vincent Noiraeux, Elias M. Puchner

**Affiliations:** School of Physics and Astronomy, University of Minnesota, Twin Cities, 55455, United States; Department of Chemistry, University of Minnesota, Twin Cities, 55455, United States

**Keywords:** PALM, SMLM, STORM, Super-resolution, Silicon Rhodamine, Oxygen Rhodamine, Photoconversion, Photoactivation, Photoswitching, Multiplexing, JF dyes, JFX650

## Abstract

The rhodamine motif has been modified in myriad ways to produce probes with specific fluorescent and chemical properties optimal for a variety of microscopy experiments. Recently, far-red emitting silicon rhodamines have become popular in single-molecule localization microscopy (SMLM), since these dyes are membrane-permeable and can be used alongside red fluorophores for two-color imaging. While this has expanded multi-color SMLM imaging capabilities, we demonstrate that silicon rhodamines can create previously unreported photoproducts with significantly blueshifted emissions, which appear as bright single-molecule crosstalk in the red emission channel. We show that this fluorescence is caused by the replacement of the central silicon group with oxygen after 640 nm illumination, turning far-red silicon rhodamines (JFX650, JF669, etc.) into their red oxygen rhodamine counterparts (JFX554, JF571, etc.). While this blueshifted population can cause artifacts in two-color SMLM data, we demonstrate up to 16-fold reduction in crosstalk using oxygen-scavenging systems. We also leverage this far-red photoconversion to demonstrate UV-free photoactivated localization microscopy (PALM) without the need for additives, and with 5-fold higher efficiency than the Cy5 to Cy3 conversion. Finally, we demonstrate multiplexed pseudo two-color PALM in a single emission channel by separating localizations by their photo-activation wavelengths instead of their emission wavelengths.

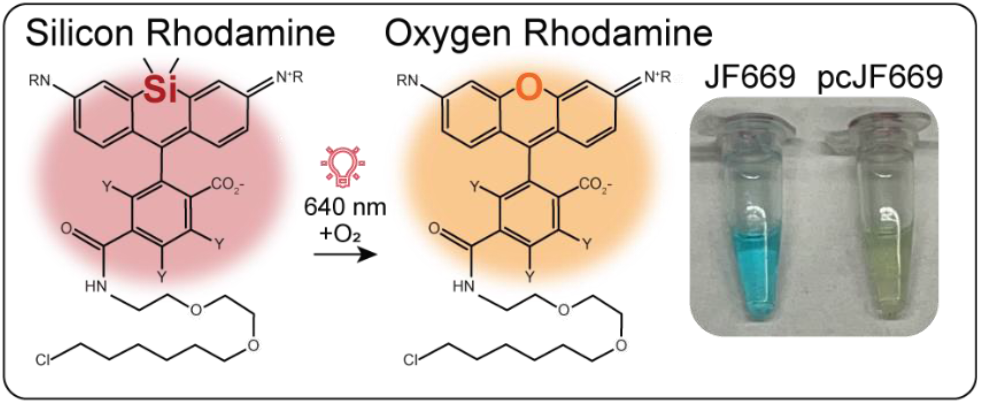

## Introduction

Single-molecule localization microscopy (SMLM) was a significant advancement for biological imaging, since it enables super-resolution microscopy of subcellular structures with ∼20 nm resolution^1^, as well as tracking of single molecules in living cells. A crucial requirement for fluorophores for SMLM is their “on’’ and “off” switching capability, which ensures each imaging frame has only one emitter within a diffraction-limited region. Many fluorophores achieve this on/off switching through an equilibrium between different fluorescent and non-fluorescent states, causing spatiotemporally well-separated bursts of fluorescence^2,3^. In contrast, photoactivatable and photo-switchable (PA/PS) fluorophores, used for photoactivated localization microscopy (PALM), change their fluorescent state through photophysical and photochemical reactions^4–6^ (Figure 1). These fluorophores are often preferred for SMLM experiments, since the emitter density can be externally controlled by the photoconversion laser intensity and does not rely on the balance of chemical equilibria that often require toxic oxidizing and reducing agents^3^. In addition, irreversible PA/PS fluorophores, which do not exhibit intrinsic and repetitive on/off cycles, enable quantification of molecule numbers in protein complexes or organelles^7–9^.

**Figure 1.**
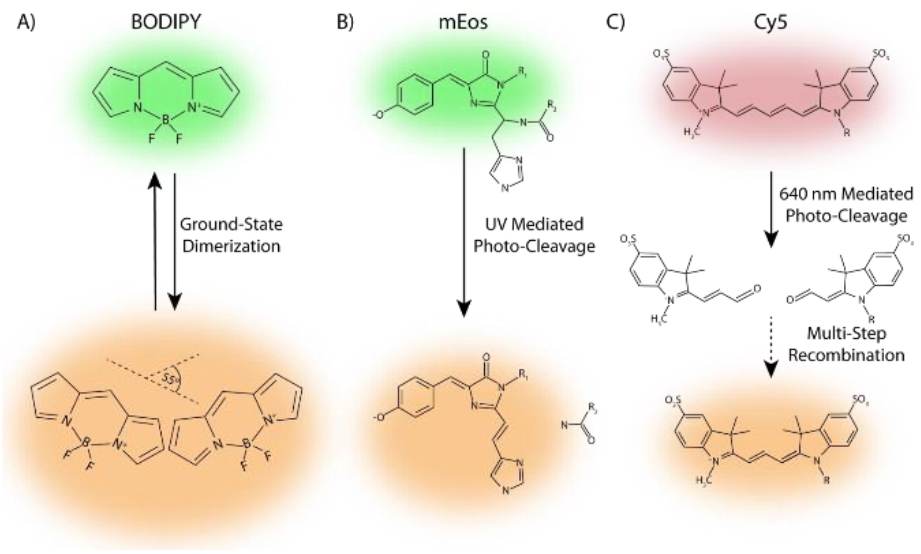
Schematics of different photoswitching and photoconversion mechanisms. A) Fluorescence switching mechanism of BODIPY through the spontaneous formation and dissociation of ground state dimers. These ground-state dimers form when two orbitals are within 3.8 Å at an angle of 55o, causing a redshift in fluorescence due to the increase in the size of the delocalized electron system across the monomers. B) Photoconversion mechanism of mEos through UV-mediated photo-cleavage, which extends the delocalized electron system, resulting in redshifted fluorescence. This general mechanism is responsible for the photoconversion of most dyes used in SMLM10,11, as well as the irreversible photoactivation of most fluorescent proteins like PAmCherry and mMaple5,6. C) Photoconversion mechanism of Cy5. 640 nm excitation causes oxidative photo-cleavage of the fluorophore, destroying fluorescence. Various recombination steps illustrated in a 2021 paper12, result in a shorter chromophore with blueshifted emission (cy3). Cy dyes are not membrane permeable and show low conversion efficiency, which peaks at 2.5%.

Silicon rhodamines have recently gained popularity for two-color SMLM since their >650 nm emissions have minimal spectral overlap with commonly used PA/PS fluorophores, like mEos and mMaple, which have emission peaks between 550 nm and 600 nm^5,6,13^. In particular, the Janelia Fluor (JF) silicon rhodamines have become increasingly popular, since they have been synthesized as conjugates for common enzymatic tags, such as HaloTag and SNAP-tag, and have superior brightness and photostability compared to other fluorescent probes^14–16^. JF dyes are also membrane-permeable and available in photoswitchable, intrinsically blinking, or fluorogenic variants^14–16^ throughout the visible spectrum, making them well-suited for multi-color SMLM imaging in both live and fixed cells. These properties, as well as their growing commercial availability, have resulted in the widespread use of JF dyes for different SMLM applications in various organisms.^17,18^

Here, we report and characterize an unexpected light-induced blueshift (∼100 nm) in fluorescence of JF silicon rhodamines, causing single-molecule fluorescence in the red emission channel. We demonstrate that this fluorescence is caused by the photoconversion of silicon rhodamines to their oxygen rhodamine counterparts upon 640 nm excitation. Oxygen rhodamines share most of the desired properties of silicon rhodamines, and are therefore also frequently used for SMLM in the red emission channel^9,19,20^. This unexpected conversion of silicon rhodamines is problematic for two-color SMLM since it causes single-molecule crosstalk in the red emission channel. Therefore, false correlations may exist in two-color data that have gone unreported thus far. We show that oxygen scavengers can be used to mitigate this conversion, greatly reducing the crosstalk from silicon rhodamines, and enabling reliable two-color SMLM experiments with minimal artifacts. While it may be beneficial in some studies to mitigate the crosstalk caused by this conversion, we also leverage this conversion for UV- and additive-free PALM in living cells. The large majority of PA/PS fluorophores rely on a relatively high intensity of UV light that is phototoxic to living cells^21^ and can cause autofluorescence artifacts in some model systems^22,23^. Additionally some pathways, like those related to DNA damage or reactive oxygen species (ROS), are currently difficult to study with PALM in living cells since the UV light required for PALM fluorophores damages DNA^24^ and creates ROS^25–27^. Among the few dye systems exhibiting UV-free photoconversion, to our knowledge, Cy5 is the only SMLM dye exhibiting additive-free and 640 nm mediated photoconversion^12^ (Figure 1). However, Cy5 cannot be used in live cells as it is not membrane-permeable, and shows low conversion efficiency only up to 2.5% in optimal conditions^12^. Styryl-coumarins conjugated to aromatic singlet oxygen reactive moieties (ASORMs) can be activated by 488 nm light without additives, and show a 68 nm blueshift of fluorescence^28^. However, the photoconverted excitation of these dyes peaks around 400 nm, causing similar complications UV-activated PALM fluorophores. In contrast, our reported photoconversion of JF silicon to oxygen rhodamines can be used for UV-free PALM in living cells, since they are membrane permeable, do not require additives or UV light, and have a conversion efficiency of up to 11.8% ± 1.8%, as shown in this work. Since silicon rhodamines can be converted with 640 nm light, while mEos2 is only activated by 405 nm light, we also utilize this difference in activation wavelengths to obtain pseudo-two-color PALM images in the same emission channel. Separating fluorescent species by their photoactivation instead of emission wavelengths removes the need for a second emission channel, simplifying the experimental setup. This imaging modality can increase spatiotemporal precision of two-color SMLM data by removing the need for imperfect transformations between emission channels and reducing pixel readout time by halving the number of pixels needed for two-color imaging.

## Results

### Red Light Converts Silicon Rhodamines to a Blueshifted State

We developed an in vitro system to study the unexpected red fluorescence from silicon rhodamines in a well-controlled chemical environment. To immobilize the dyes for accurate quantification, we utilized a cell-free transcription-translation (TXTL)^29^ system expressing HaloTag fused to the mono-streptavidin domain (MSA), which irreversibly links bound JF dyes to the biotinylated surface of coverslips (Figure 2A). Using this system, we immobilized and imaged JFX650 with alternating 640 nm and 561 nm excitation to probe their red and far-red populations. Consistent with the emission spectrum of JFX650 control samples (Figure 2B,) we initially observed significant far-red fluorescence, with only a minimal background signal in the red emission channel. However, after repeated exposure to 640 nm light throughout imaging, we observed up to 20-fold increase in fluorescence intensity in the red emission channel. A similar photoconversion was also observed for the other silicon rhodamines JF669 and JF635 (Figure S1A). The oxygen counterpart for JF635 was not available for comparison but has been well characterized in previous studies^25–27^. While405 nm and 640 nm light caused the largest activation, a small yet significant amount of activation was also observed with 561 nm light (Figure S1B).

**Figure 2.**
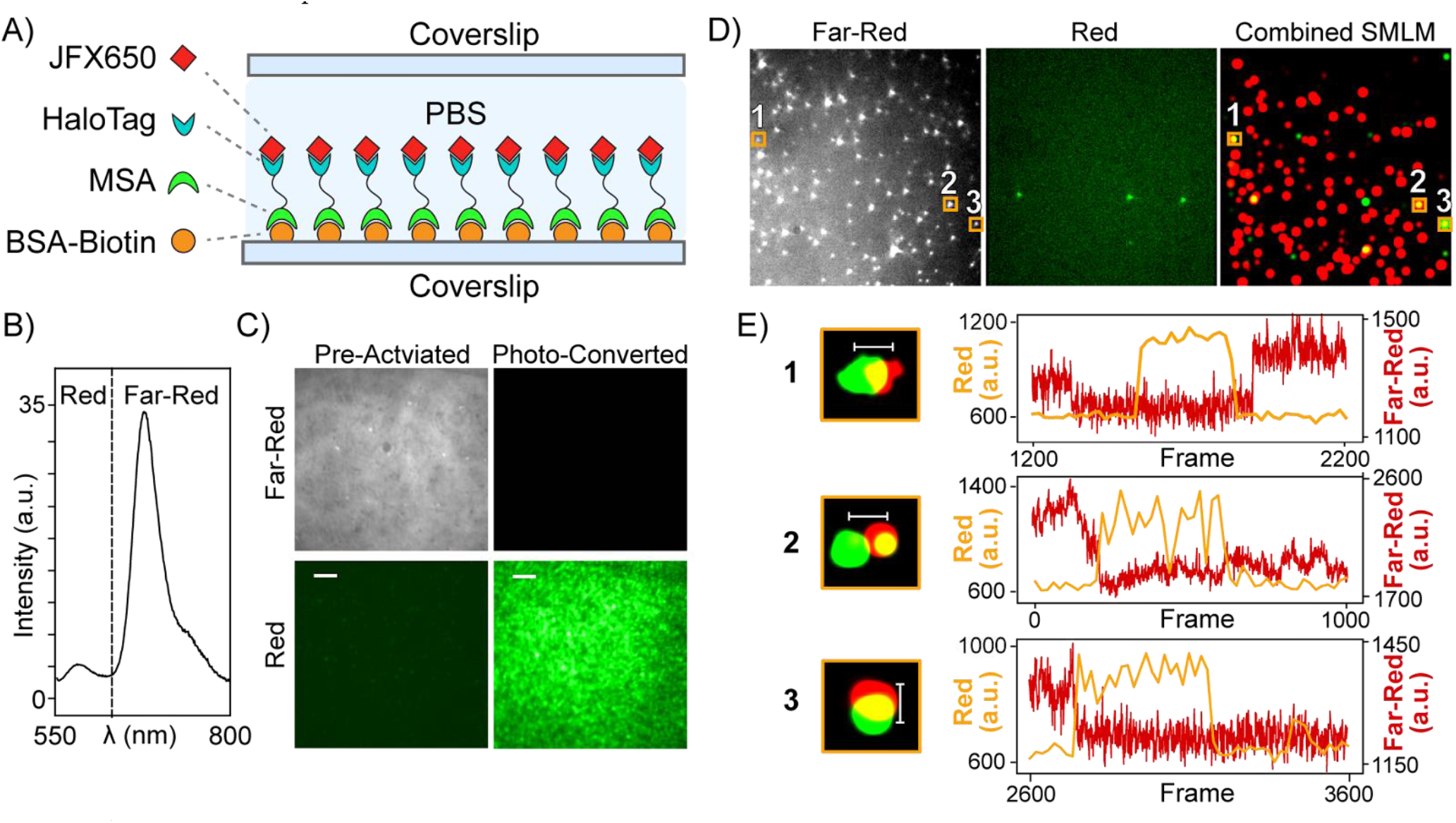
A) Schematic of the *in vitro* setup to characterize fluorophores. The TXTL system was used to express HaloTag fused to a mono-streptavidin domain (MSA). The HaloTag protein binds JFX650 while the streptavidin domain binds to the biotinylated-bovine serum albumin (BSA-Biotin) on the plasma-cleaned coverslips. B) Emission spectrum of JFX650 under 520 nm excitation with a dotted line indicating the split between the red and far-red emission channels of the used microscope. C) Initially JFX650 only exhibits far-red fluorescence and no significant red signal. After illumination with 640 nm light for ten seconds, significant and unexpected fluorescence is detected in the red channel with 561 nm excitation. D) The in vitro system was sparsely labeled with spatially well-separated JFX650 as seen in the raw image of the far-red emission channel (left). The red-channel fluorescence of pc-JFX650 (middle) shows significant co-localization with the JFX650 signal as is seen in the merged single-molecule image (right). 169 ± 11 localization clusters from single fluorophores are detected in the far-red emission channel and 20 ± 3 localization clusters are detected in the red emission channel, resulting in a conversion efficiency of 11.8% ± 1.8%. E) Left: Magnified regions of three selected molecules showing co-localization. Scale bar 100 nm. Right: The fluorescence intensity traces of the magnified regions show single molecule transitions with most molecules exhibiting a stepwise disappearance of far-red fluorescence before showing a stepwise increase in red fluorescence, and rarely the reverse can be seen as in trace 1.

These results suggest that silicon rhodamines undergo a light-induced chemical or conformational change, which switches their fluorescence to a blueshifted state. 405 nm photoconversion is unsurprising, as the majority of PA/PS fluorophores, like mEos, are activated through UV-induced photocleavage^11^. This conversion is common due to the relatively high energy of these photons, despite the low absorption at these wavelengths for some photoactivatable fluorophores^4^. While the other wavelengths show minimal silicon rhodamine conversion, 640 nm light causes both significant conventional fluorescence excitation and photoconversion to the blueshifted state, suggesting that the excited state may play an important role in the 640 nm mediated photoconversion.

Shifted excitation and emission spectra can result from dye-dye interactions in other systems like BODIPY, which form redshifted ground state dimers suitable for SMLM^30–33^(see also Figure 1). To rule out the possibility of such dye-dye interactions, we immobilized JFX650 on coverslips at a sufficiently low density such that single emitters in the far-red channel were separated by distances much larger than the diffraction limit (Figure 2E left). The red channel also showed well-separated fluorescent bursts that appeared for many consecutive frames (Figure 2E middle) and co-localized well with the far-red localizations (Figure 2E right). Magnified regions show red and far-red co-localization within the localization precision of the single molecule signals. Importantly, the intensity vs. time traces of the selected regions with co-localization show a stepwise loss of far-red fluorescence before a stepwise emergence of red fluorescence. The stepwise transitions in sparsely labeled samples indicate that individual JFX650 molecules change their fluorescent state without interacting with other dye molecules. To determine the conversion efficiency, localizations in each channel were clustered to determine the number of molecules in each state. These experiments showed 169 ± 11 molecules in the far-red emission channel, and 20 ± 3 in the red emission channel, resulting in a conversion efficiency of 11.8% ± 1.8%. While we observed a significant number of traces showing conversion from far-red to red fluorescence, we also observed a few instances of the reverse reaction, suggesting this process may be reversible (Figure 2F, 1). However, due to the low labeling density and limited conversion efficiency, only limited statistics of the reverse transition was obtained.

### Replacement of the Central Silicon Group with Oxygen Causes Blue-shifted Emission

To determine the identity of the red photoproducts, we first measured the emission spectra of different silicon rhodamines during bulk photoconversion with 640 nm light (Figure 3A,B). Since the power density of the laser used for bulk photoconversion is 22,000 times lower compared to the microscopy experiments, instead of seconds, it takes days in bulk to deliver the equivalent amount of 640 nm energy to each dye molecule. JFX650 exhibited a decrease in its emission peak at 667 nm while at the same time, a distinct blueshifted emission peak emerged from photoconverted JFX650 (pc-JFX650) at 576 nm. After 72 h of photoconversion, most of the fluorescence signal was due to pc-JFX650, and the 576 nm peak matched that of the oxygen rhodamine variant, JFX554. Likewise, the photoconversion of JF669 resulted in the emergence of a blueshifted emission peak at 584 nm, which matched its oxygen counterpart JF571. However, an additional gradual 22 nm photoblueing occurred over the course of 72 hours. During photoconversion, numerous compounds are generated, and while most are nonfluorescent, slight modifications to the dye structure can result in small spectral shifts. ∼20 nm blueshift has been previously reported for similar JF dyes, and results from a well-characterized breakdown of the same side chains that are also present in JF669^34^. When JF571 was exposed to the same conditions as JF669, a similar photoblueing was observed, and the photoblued JF571 (pb-JF571) peak matched that of pcJF669 at 562 nm. JFX dyes differ in the design of the side chains and showed significantly less photoblueing in our experiments, which is likely due to the increased stability of JFX side chains^14^ (Figure 3B). While this sidechain breakdown explains the minor broadening and shifting of emission peaks, it cannot explain the single-molecule fluorescence in the red emission channel, which is caused by a population that exhibits a much larger blueshift (∼100 nm) from a different reaction (Figure 3 A, B).

**Figure 3.**
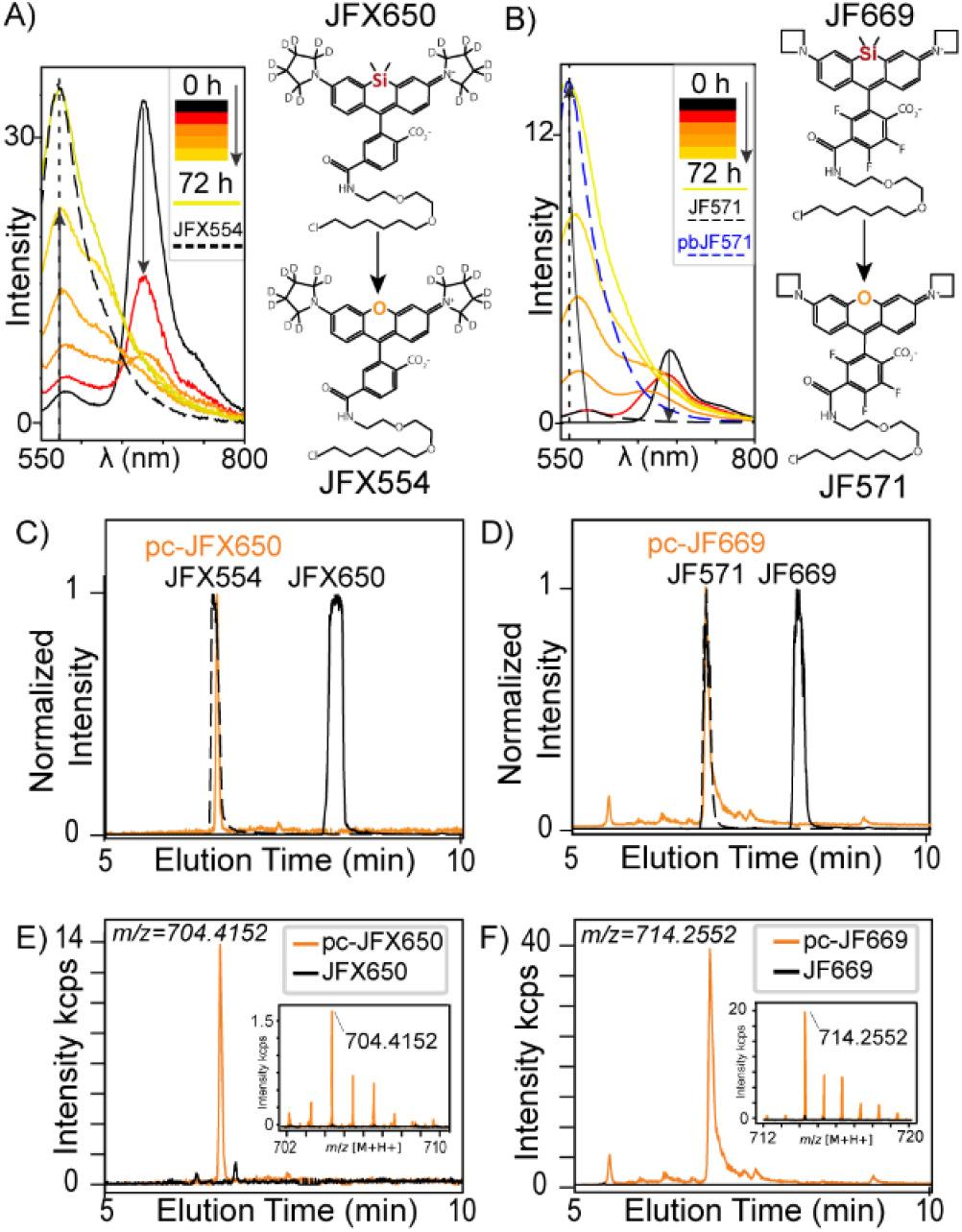
A) Emission spectra of JFX650 after illumination with a 640 nm laser for various times up to 72 h. The legend indicates the illumination time from red to yellow, of t=0,1 h, 5 h, 24 h, 48 h, and 72 h respectively. A continuous decrease in farred fluorescence as well as an increase in red fluorescence is observed, which peaks within 1 nm of the emission peak of JFX554 (dashed), the oxygen counterpart of JFX650. Right: Chemical structures of JFX650 and the corresponding oxygen rhodamine JFX554. B) Emission spectra of JF669 after illumination with a 640 nm laser for various times up to 72 h. A similar trend as for JFX650 is observed, however, JF669 and JF571 are slightly photoblued during activation. After 72 h, the emission peak of photoconverted JF669 matches the emission peak from JF571, which also has been photoblued for 72 h by exposure to 640 nm light. C) UHPLC\QTOF-MS results from JFX650, JFX554, and pc-JFX650. The normalized Selected Ion Chromatograms (SIC) show JFX554 and JFX650 elution at significantly different times, while, pc-JFX650 elutes at the same time as JFX554. D) The same experiment repeated for JF669 and JF571 show similar results, suggesting oxygen replaces the silicon group in both dyes under 640 nm photoconversion. E) SIC of m/z=704.4152 (+/-0.0025 Da) detect JFX554 only in the photoconverted sample. Mass spectra of the JFX650 control and pc-JFX650 are inset. F) SIC of m/z=714.2552 detects JF571 only in the photoconverted sample. Mass spectra of JF669 control and pc-JF669 are inset.

While the spectral analysis suggests a possible photoconversion of silicon rhodamines to their oxygen counterparts, it is not sufficient to determine the chemical identity of the photoconversion products. To verify that the oxygen counterparts are generated during photoconversion, we utilized UHPLC/QTOF-MS, which is sufficiently accurate to determine elemental composition^35^. In selected ion chromatograms (SICs) corresponding to JFX650, JFX554 and pc-JFX650, pure oxygen rhodamine control samples eluted at the same time as the suspected oxygen rhodamines in the photo-converted samples. In contrast, the silicon rhodamine control eluted much later (Figure 3C). Similar results were observed for JF669, JF571, and pc-JF669. Additionally, since the control samples showed no significant peak in the SICs corresponding to the oxygen rhodamines (Figure 3 E, F), the suspected oxygen rhodamine was only present after photoconversion. This is corroborated by the mass spectra of the photoproducts shown in the insets of Figure 3E and Figure 3F. These mass spectra match those of the oxygen rhodamine controls with high mass accuracy (Figure S2, S3). Additionally, some of the suspected photoblued compounds of JF669 and JF571 were also detected in the photoconverted samples (Figure S4, S5)

In summary, LC/MS results validate the chemical composition while the emission spectra and chromatography results are both consistent with those of the oxygen rhodamines. We therefore conclude that the photoconversion of silicon rhodamines in response to 640 nm illumination produces oxygen rhodamines, which are responsible for the single-molecule fluorescence in the red emission channel. The three silicon rhodamines tested here (JF669, JFX650, and JF635) share the same HaloTag ligand and central silicon rhodamine motif but differ in their side groups and fluorinations. These results suggest central group replacement occurs regardless of modification to these groups, and likely occurs in many silicon rhodamine variants.

### Oxygen Scavenger Inhibits Crosstalk from Photoconversion

The generation of blueshifted oxygen rhodamines creates unwanted and unexpected crosstalk in the red emission channel during two-color SMLM experiments. If unaccounted for, a significant number of localizations could be associated with the incorrect label, creating spurious correlations. Since this photoconversion requires oxygen, we utilized the glucose oxidase and catalase (GOC) oxygen scavenging system to reduce molecular oxygen in solution and to inhibit photoconversion. Using our *in vitro* system (Figure 2A) we again alternated 640 nm and 561 nm light to convert and measure the amount of JFX554 created during imaging (Figure 4A). The control sample without oxygen scavengers showed higher photoconversion and photobleaching compared to the samples with GOC, which is expected as both processes are oxygen-dependent. At the highest tested concentration of glucose oxidase (66 U/mL) and catalase (6.6 U/ML) in a PBS solution containing glucose (4 g/L), a maximum 16-fold reduction in peak red fluorescence was seen, confirming the oxygen dependence of the photoconversion.

**Figure 4.**
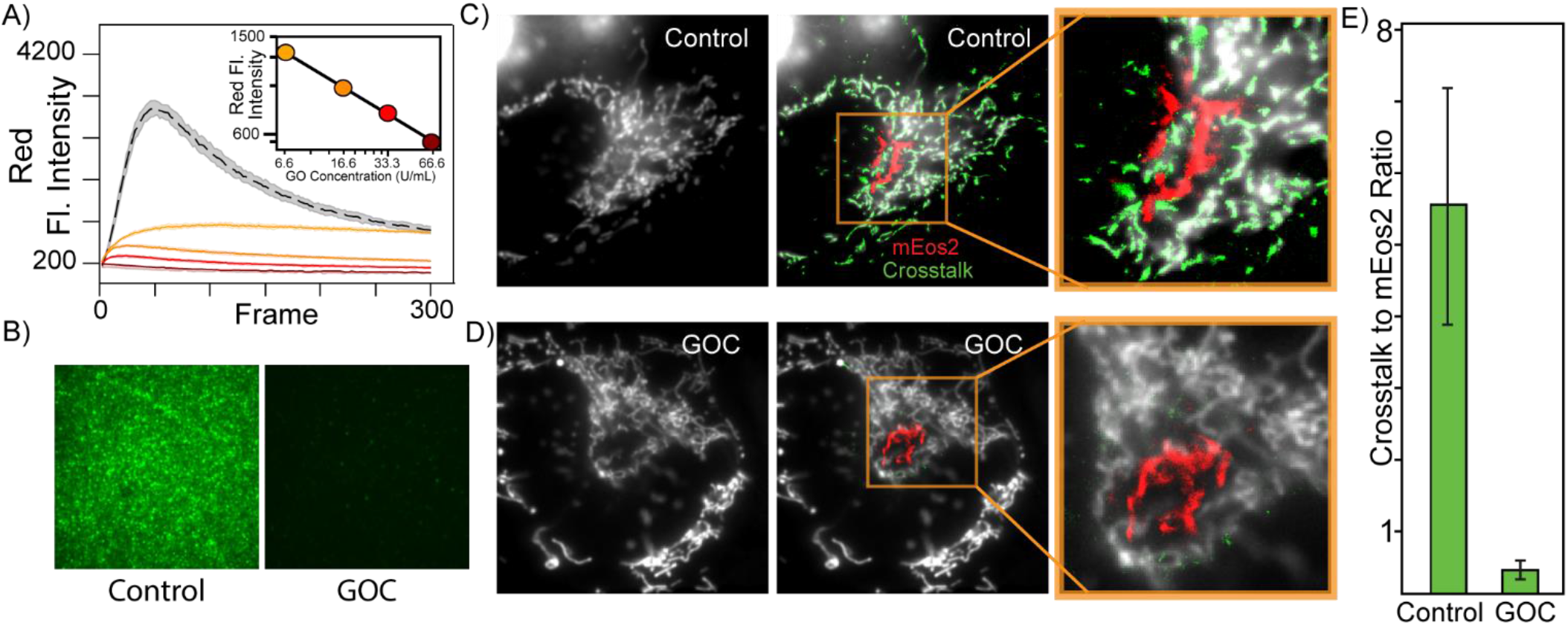
A) JFX650 was immobilized on coverslips and imaged with alternating 640 nm and 561 nm light to measure photoconversion rates with increasing oxygen scavenger concentrations as indicated. Control samples without GOC in black show the most red fluorescence activation as well as the highest rate of bleaching, demonstrating the oxygen dependence of both processes. Inset is a log-log plot of peak red fluorescence intensity as a function of glucose oxidase concentration. B) Red fluorescence images of JFX650 samples with and without GOC after 10 seconds of photoconversion by 640 nm light. C) HeLa cells were transiently transfected with mEos2 targeted to Golgi, and HaloTag, labeled with JF669, targeted to the inner mitochondrial membrane. The conventional far-red fluorescence image average (white, left) shows JF669 on the mitochondria. PALM imaging using the 640 nm, 561 nm, and 405 nm laser shutter sequence (Figure S9) to demonstrate the potential for single-molecule crosstalk in the red emission channel. Localizations in the red channel were rendered on top of the conventional image (middle), with localizations that colocalize with the conventional signal (JF669) being colored green, and the rest being colored red (mEos2). 7x zoom (right). D) The same system was imaged with the inclusion of 66.6U/mL of Glucose Oxidase and 6.6U/mL of Catalase, showing minimal red single-molecule signal from the mitochondria, while maintaining the far-red fluorescence signal. E) The ratio of unwanted mitochondrial localizations to intended mEos2 localizations showed a 16-fold reduction in crosstalk with the inclusion of GOC.

To demonstrate the potential for crosstalk in two-color SMLM as well as its suppression, we imaged two transiently expressed constructs in live HeLa cells, targeting mEos2 to the surface of the Golgi and HaloTag-JF669 to the inner mitochondrial membrane (Figure 4B). To image mEos2 at the Golgi, a typical PALM laser shutter sequence was used, consisting of 640 nm, 561 nm, and 405 nm illumination (Figure S6). Localizations were separated based on the conventional far-red fluorescence signal from JF669 on the mitochondria and the well-defined morphology of the two labeled organelles. Localizations that co-localized with the far-red fluorescence signal were identified as crosstalk from the JF669-labeled mitochondria (Figure 4B, green) and the remaining localization were identified as mEos2 on the Golgi (Figure 4B, red). While the large amount of crosstalk and obvious difference in morphology make it easy to identify crosstalk in these data, in systems with lower expression and less obvious morphological differences, photo-converted silicon rhodamine localizations could easily remain unnoticed and be misidentified as mEos2 or other red fluorophores. This crosstalk is especially misleading when the two labeled proteins are hypothesized to be highly correlated or co-localized. To mitigate this crosstalk, we used the highest GOC concentration tested, which reduced the number of localizations caused by the pc-JF669, without significantly reducing the far-red signal from JF669 on the mitochondria or from mEos2 on the Golgi (Figure 4C). The ratio of crosstalk to mEos2 localizations showed a 16-fold reduction in samples containing GOC (Figure 4D), which agrees with *in vitro* experiments (Figure 4A). For this reason, we recommend that some form of oxygen scavenger be included in experiments that require red and far-red imaging of silicon rhodamines.

### Silicon Rhodamines Enable UV- and Additive-Free PALM

Although the photoconversion of silicon to oxygen rhodamines can cause crosstalk if not accounted for, with knowledge of the factors affecting photoconversion, this blueshifted population can be utilized for new imaging modalities. One promising application of this conversion is UV-free and additive-free PALM, which avoids the harmful effects of phototoxicity and toxic chemicals in live-cell experiments. To demonstrate this application, we expressed HaloTag-TOM20, targeting JF669 to the outer mitochondrial membrane. By alternating 640 nm activation light and 561 nm excitation light (Figure S7A), we obtained conventional fluorescence images in the far-red (Figure 5A,) and super-resolution images in the red channel without any additives or UV light (Figure S7B). The photoconversion created enough localizations to resolve the sub-diffraction-limited structure of the mitochondria, which can be clearly visualized in the magnified images (Figure 5C). Not only yielded this imaging enough localizations to resolve these sub-200 nm structures, but the generated oxygen rhodamine was also remarkably photostable. In these data, pc-JF669 produced a large number of single-molecule traces longer than 0.25 s (Figure 5D) with many being longer than one second. These on-times are significantly longer compared to mEos2, which showed no on-times longer than 0.25 s. pc-JF669 therefore enables a more accurate determination of diffusion coefficients in single molecule tracking, and a higher localization precision per fluorophore due to the higher photon counts (Figure S8). For these reasons, silicon rhodamines can be preferable compared to mEos2 for tracking experiments, especially when studying stress-dependent pathways that may be altered by high intensity UV illumination.

**Figure 5.**
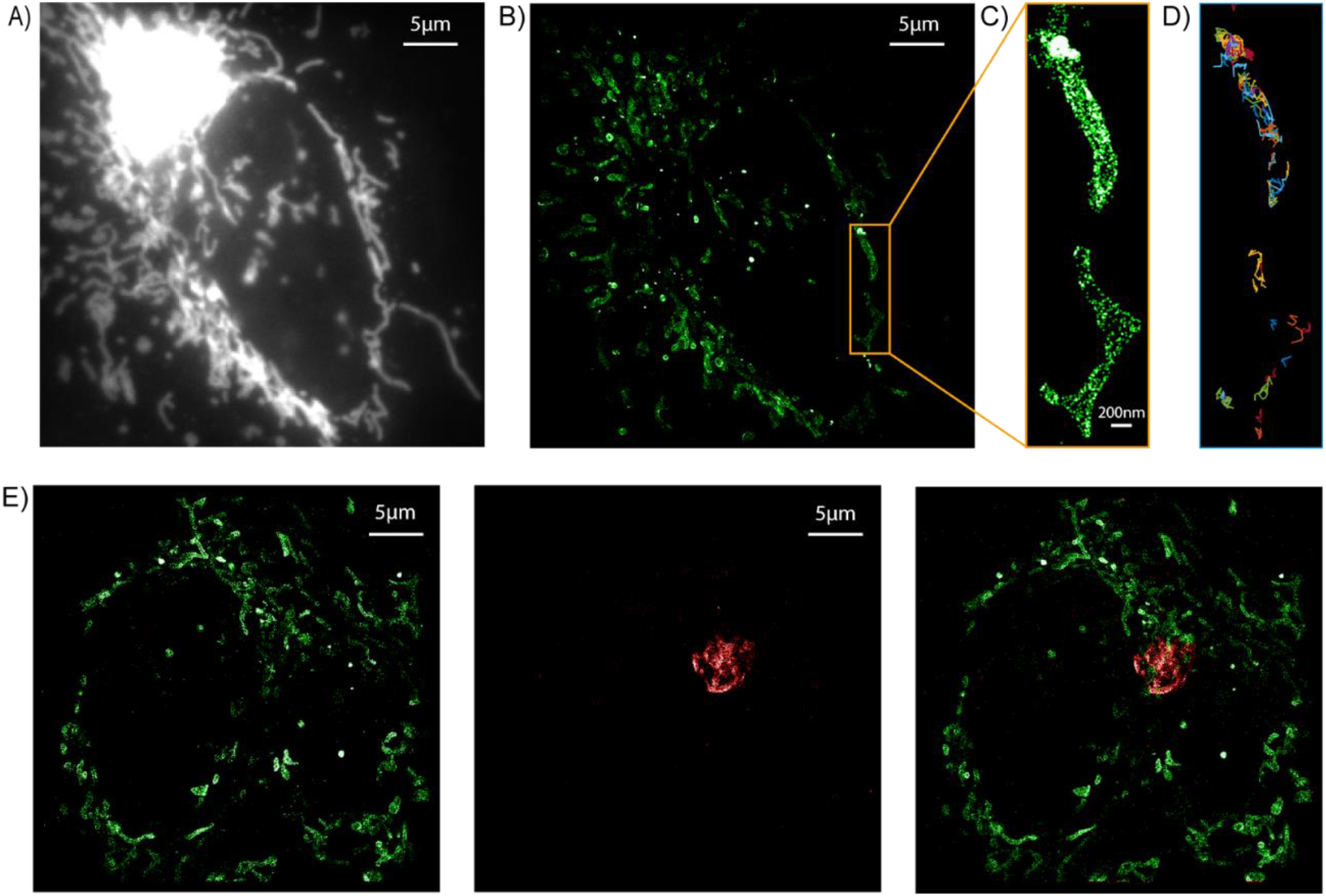
A) Conventional far-red fluorescence image of JF669 labeling TOM20 on the outer mitochondrial membrane. B) SMLM image of pc-JF669 acquired without UV or additives shows sufficient localization density to resolve diffraction-limited structures of the mitochondria. C) Zoomed image D) Single-molecule traces of pc-JF669 with on-times > 0.25 s are significantly longer than from mEos2. E) HeLa cells were transiently transfected with HaloTag on the outer mitochondrial membrane (green) and mEos2 on the Golgi (red) to obtain a pseudo-two-color SMLM image in the same emission channel. First, the mitochondria were imaged (left) by alternating 640 nm activation light with 561 nm excitation light. After JF669 was imaged, a typical PALM shutter sequence was used with UV to image mEos2 on the Golgi (middle), showing almost no crosstalk between the two signals (right).

### Silicon Rhodamine Photoconversion Enables Pseudo Two-color PALM in a Single Emission Channel

Since mEos2 is not activated or excited by 640 nm light, silicon rhodamines can first be imaged and bleached without UV light before photoactivating and imaging mEos2. In this way, pseudo-two-color PALM images can be created in the same emission channel by separating fluorescent labels by their activation wavelengths instead of their emission wavelengths. To demonstrate this multiplexed imaging modality, we sequentially imaged the mitochondria, labeled by TOM20-HaloTag (JF669), and mEos2-B4GALT1, which labeled the Golgi. By first using alternating 640 nm and 561 nm illumination (Figure S7A) all the silicon rhodamines on the outer mitochondrial membrane were imaged (Figure 5E left,). Next, alternating 405 nm and 561 nm light (Figure S7B) was used to image mEos2 on the Golgi (Figure 5E middle). The images show virtually no crosstalk and can be merged to reveal the relationship between the Golgi and Mitochondria in the same emission channel.

This pseudo-two-color imaging has multiple benefits over the traditional method of separating fluorophores by emission wavelengths. Microscopes with two emission channels require camera chips that are at least twice as large to image the same object in two different colors and are therefore more expensive. Additionally, splitting the signal based on emission wavelength requires additional lenses, dichroic mirrors, and filters in the emission path of the microscope, adding to the expense and complexity of the experimental setup. These optics also result in a loss of photons and introduce linear and nonlinear distortions to the signal. To precisely superimpose two-color SMLM images and to account for these distortions, a transformation between the two emission channels^23^ needs to be determined based on two-color SMLM images of a calibration standard. However, this transformation is never perfect and results in a loss of resolution equal to the localization uncertainty of the calibration standard, and in practice can result in a 10-60 nm loss in resolution due to uncertainties in the transformation (Figure S9). Compared to traditional two-color PALM, this new imaging approach can therefore provide increased resolution, while simplifying experimental setup and reducing costs.

## Conclusion

We reported and characterized a new photoconversion of silicon rhodamines, which exhibited unexpected red single-molecule fluorescence after 640 nm photoactivation. The emission spectra, LC/MS results, and singlemolecule fluorescence traces show that silicon rhodamines convert to oxygen rhodamines after 640 nm illumination, causing single-molecule crosstalk in the red emission channel. Since the conversion efficiency can be as high as 11.8%, it is possible that previous studies utilizing silicon rhodamines for two-color imaging may have overestimated the correlation between the two labeled populations. We demonstrate that the glucose oxidase and catalase (GOC) oxygen scavenging system can be leveraged to reduce the conversion rate by up to 16-fold, showing clear oxygen dependence as well as GOC’s efficacy in mitigating unwanted crosstalk. The detailed photochemical reaction of this conversion has yet to be determined, and further studies are needed to clarify whether the reaction requires molecular oxygen or other ROS. Regardless, this conversion shows promise for use as a single-molecule fluorescent sensor of the oxygen species involved in the reaction. Finally, we utilized this new conversion to demonstrate UV- and additive-free PALM as well as pseudo two-color PALM in a single emission channel.

## ASSOCIATED CONTENT

Additional experimental details, materials and methods, UHPLC/QTOF-MS results, DNA used, background measurements, and localization information is provided in the supporting information. Raw Videos also included.

## AUTHOR INFORMATION

## Author Contributions

The manuscript was written through contributions of all authors. / All authors have given approval to the final version of the manuscript.

## Notes

The authors declare no competing financial interest.

## ACKNOWLEDGMENT

Special thanks to the Luke Lavis lab, which provided the dyes as part of the open chemistry initiative, and also to Ziane Izri for helpful conversations.

## ABBREVIATIONS

PALM: Photoactivated Localization Microscopy
SMLM: Single-Molecule Localization Microscopy
Cy: Cyanine
JF: Janelia Fluor
TXTL: Transcription-Translation
UV: Ultraviolet.
PC: Photoconverted
PA: Photoactivateable
PS: Photoswitchable
ASORM: aromatic singlet oxygen reactive moieties.
ROS: Reactive Oxygen Species
GO: Glucose Oxidase
C: catalase
GOC: Glucose Oxidase and catalase
SIC: Selected Ion Chromatogram,

