## Supplemental Info for "Red Light Mediated Photo-Conversion of Silicon Rhodamines to Oxygen Rhodamines for Single-Molecule Microscopy"

### Table of Contents

1) Camera setting and calibration
2) Conversion experiments
3) UHPLC/QTOF-MS experiments and analysis
4) Spectral measurements
5) Microscope and lasers
6) PALM crosstalk imaging
7) UV-free and additive-free PALM
8) Emission channel transformation
9) Pseudo two-Color PALM in the same emission channel
10) TXTL
11) *In vitro* sample preparation, imaging and analysis

#### 1. Camera settings and calibration

All movies (conventional and single molecule) were recorded at a frame rate of 20 Hz on an electron-multiplying CCD camera (Ixon 89 Ultra DU-897U; Andor), which was cooled down to −70 °C and the amplifying EMCCD gain was set to 30 for single-molecule measurements and 0 for fluorescence Intensity measurements. All imaging data was recorded at 20 Hz frame rate. The EMCCD camera calibration was performed to determine the gain (e/ADU) and the camera offset. The camera pixels were exposed to varying intensities of light with an EMCCD gain of 30. The slope of the mean intensity vs. variance graph accounting for the extra noise factor *F*2 was used to calculate the gain. The calculated gain of 0.166 e/ADU matched the manufacturer's specifications and was used to convert the integrated CCD counts to photons.

#### 2. Conversion experiments

1 µM samples of JF dyes in PBS were exposed to different laser wavelengths on the microscope for 10s at 3.4 kW/cm$^2$ to determine the wavelength specificity of photoconversion. The red fluorescence was measured with 561 nm light at the same power to probe the photoconverted population.

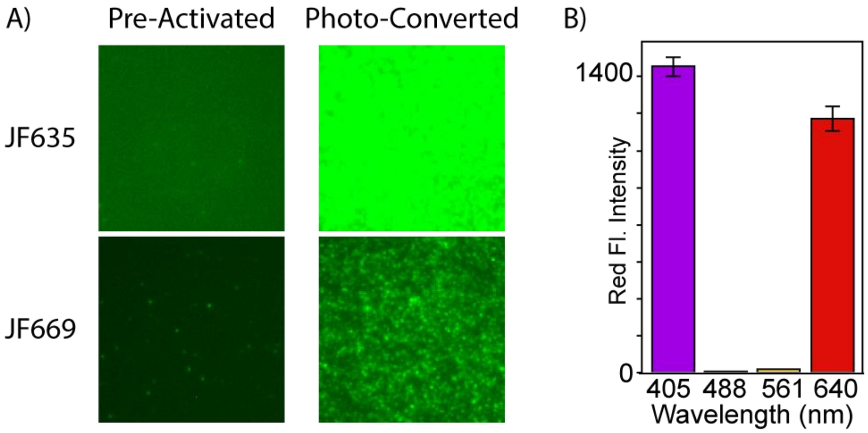


Figure S1. A) Fluorescence images of red channel fluorescence JF635 and JF669 at 1 µM before and after 10 s of Illumination at 3.4 kW/cm$^2$ with 640 nm light, showing that these dyes show a similar blueshifted conversion. B) Photoconversion at different wavelengths. Samples of JFX650 immobilized on coverslips were imaged at 1 µM in PBS. After 10 s of Illumination at 3.4 kW/cm$^2$ with 405 nm, 488 nm, 561 nm, or 640 nm light, the photoconverted population was probed using 561 nm light for a single frame at the same power density.

#### 3. UHPLC/QTOF-MS analysis

A Sciex Exion UHPLC coupled to a Sciex X500R quadrupole time-of-flight (Q-TOF) mass spectrometer was used for separation and accurate mass measurement of the dye samples. An Agilent XDB $C_{18}$ UHPLC $C_{18}$ 2.1 mm x 100 mm column (1.8 µm particles) at 40° C was used during the following 21 min gradient separation with A: Water containing 0.1% formic acid and B: ACN containing 0.1% formic acid, at a flow rate of 0.5 mL/min: 20% B, 0 min to 2 min; 20% B to 70% B, 2 min to 5 min; 70% B to 97% B, 5 min to 15 min; 97% B, 15 min to 17 min, 97%

B to 20% B, 17 min to 18 min, total run time 21 min. The autosampler was kept at 10° C, and the injection volume was 30 µL. Positive ionization electrospray mass spectra were collected every 0.25 s over the *m/z* range 50-1200 during the chromatographic separation detailed above. MS parameters were as follows: capillary voltage, 5500 V; temperature, 500 °C; ionization gas (1), 38 psi; ionization gas (2), 38 psi; curtain gas, 30 psi; CAD gas, 7 psi; declustering potential (DP), 50V; DP spread, 0 V; collision energy (CE), 10 V; CE spread, 0 V. All samples were 200 µL of 500 µM dye samples in DI water in 0.5 mL plastic Eppendorf tubes. Photoconverted samples were illuminated for 2 h directly in front of the 640 nm laser before UHPLC/QTOF-MS measurements.

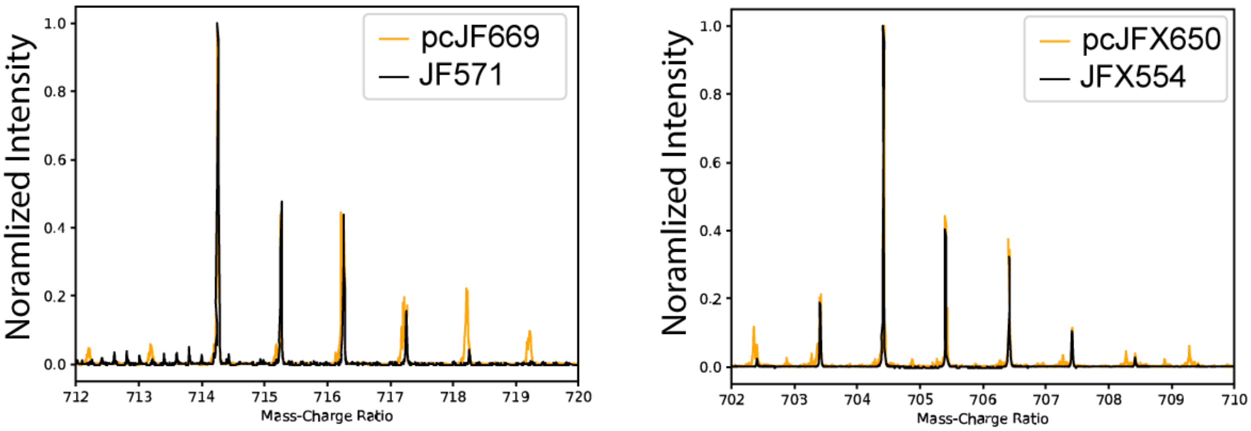


Figure S2. Selected ranges of mass spectra to verify proposed elemental composition of compounds. pcJF669 ion at *m/z* 714.2551 (theoretical *m/z* for $C_{37}H_{39}Cl_1F_3N_3O_6$, 714.2552, 0.14 ppm mass measurement error). pcJFX650 ion at *m/z* 704.4166 and JFX554 ion at *m/z* 704.4169 (theoretical *m/z* for $C_{39}H_{30}D_{16}Cl_1N_3O_6$, 704.4152, mass measurement errors 2.4 ppm and 2.0 ppm, respectively.)

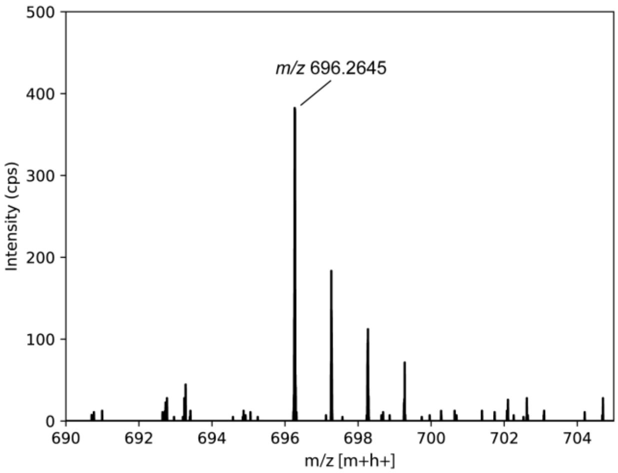

Figure S3. Selected range of mass spectra to verify proposed elemental composition of compounds. pcJF635 ion at m/z 696.2645 (theoretical *m/z* for $C_{37}H_{40}Cl_1F_2N_3O_6$, 696.26462, -0.17 ppm mass measurement error).

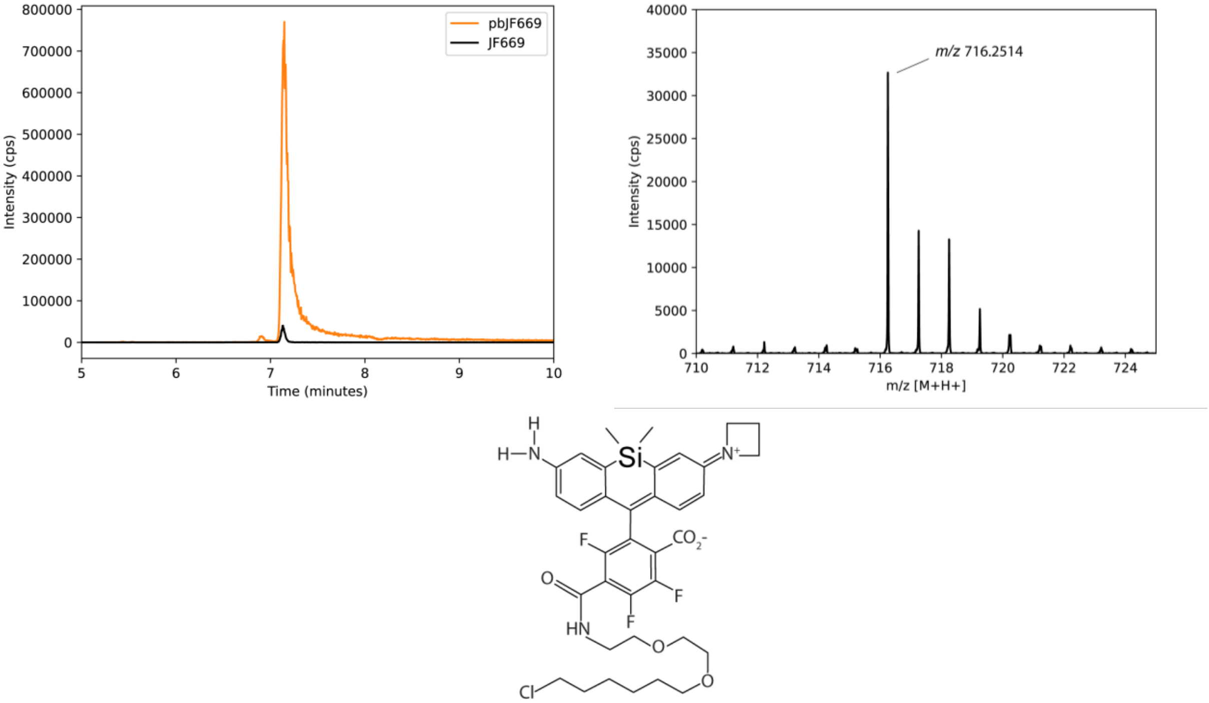


Figure S4. Selected range of normalized Selected Ion Chromatograms (SIC) and mass spectrum to verify proposed elemental composition of compounds. Photo-degradation causes the generation of many products, and some of them maintain fluorescence. Since the aim of this article is not to characterize all products, we only include evidence of one of the photoblued products[35], which is motivated by previous characterization of photoblued rhodamines[35]. SIC and MS show this compound is generated during photoconversion and may explain the slight broadening and shifting of emission peaks in JF669 samples. pb-JF669 ion at *m/z* 716.2514 (theoretical *m/z* for $C_{36}H_{41}Cl_1F_3N_3O_5Si_1$, 716.2529, -2.04 ppm mass measurement error).

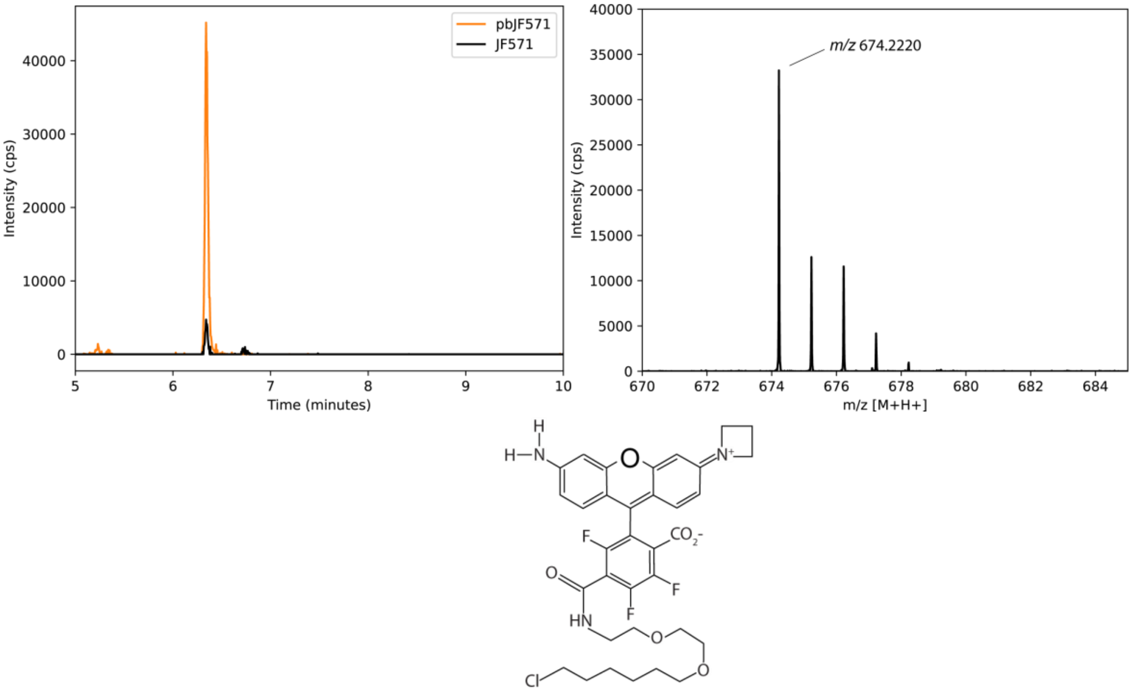


Figure S5. Selected range of normalized Selected Ion Chromatograms (SIC) and mass spectrum to verify proposed elemental composition of compounds. pb-JF571 ion at *m/z* 674.2220 (theoretical *m/z* for $C_{34}H_{35}Cl_1F_3N_3O_6$, 674.2239, -2.81 ppm mass measurement error) detected in photo-blued JF571 samples. SIC and MS show the displayed compound is generated during photoconversion and may explain the slight broadening and shifting of emission peaks in JF669 samples.

#### 4. Spectral Measurements

Each silicon rhodamine was diluted to 500µM in DI water, and 200µL was placed in 0.5 mL plastic PCR tubes (Fisher, cat. 14230200) for photoconversion. Samples were placed directly in front of the 640 nm laser and wrapped in aluminum foil to create a laser cavity and increase efficiency. At 0 h, 1 h, 5 h, 24 h, 48 h, and 72 h, 10 µL of the sample were placed in a 384 well microplate (Greiner Bio-One, cat. 781096) and placed in the Agilent BioTek Synergy Neo2 Hybrid Multimode Reader. Spectral measurements were taken at 37 °C with 520 nm +/- 20 nm excitation wavelength. The emission intensities were measured starting at 550 nm +/- 10 nm with 1 nm steps until 800 nm. Fluorescence was measured from the bottom of the well with a gain of 150. 100 measurements were averaged for each data point to generate emission spectra.

#### 5. Microscope and lasers

All microscopy experiments were performed with a Nikon Ti-E inverted microscope with a Perfect Focus System. Lasers emitting at 405 nm, 488 nm, 561 nm, and 640 nm, (OBIS-CW; Coherent) were combined using dichroic mirrors, aligned, expanded, and focused to the back focal plane of the objective (Nikon-CFI Apo 100× Oil immersion N.A 1.49). A quad band dichroic mirror (zt488/561/640rdc; Chroma) filters out excitation light and reflects emission into the emission path. For simultaneous red and far-red imaging, the emission was split by a dichroic longpass beamsplitter (FF652-Di01; Semrock) and further filtered by bandpass filters: ET610/75 (Chroma) in the red and FF731/137 (Semrock) in the far-red channel. The lasers were controlled directly by a computer through the Hal 4000 program (Xiaowei Zhuang lab, Harvard). The laser powers and shutter sequences were controlled directly by a computer through the Hal 4000 program (Xiaowei Zhuang lab, Harvard).

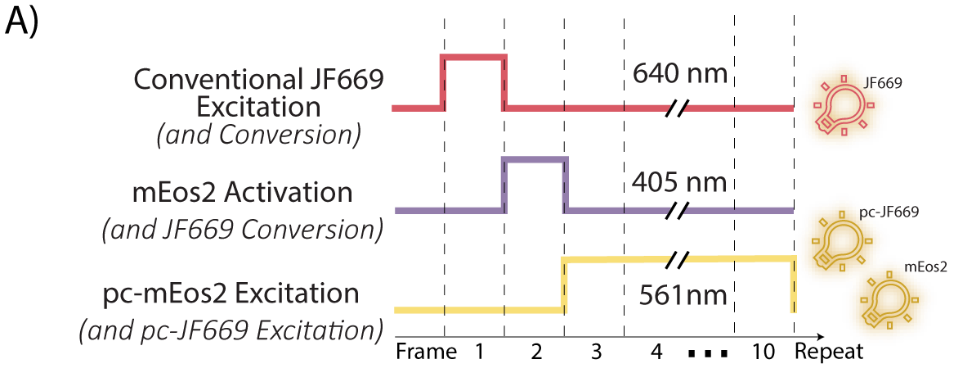


Figure S6. Typical PALM shutter sequence used to image cells containing mEos2 and JF669. This imaging has the intent to only image mEos2 in the red and JF669 in the far-red channel, but a significant number of single-molecule signals from photo-converted JF669 is produced in the red channel despite having only a single 640 nm frame per cycle.

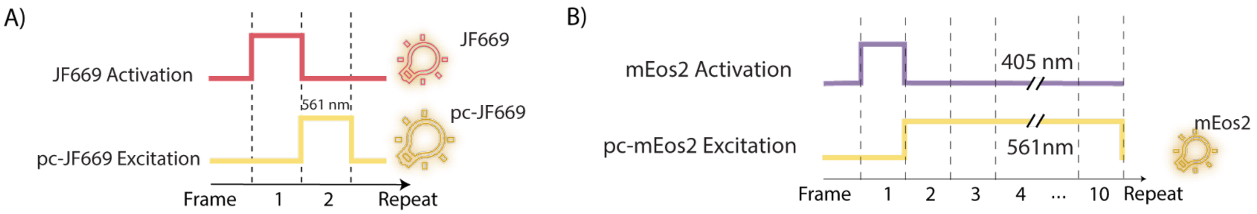


Figure S7. PALM shutter sequences used for pseudo-two-color PALM in the red emission channel. A) 640 nm light is used to activate JF669, while minimally affecting mEos2. 561 nm light excited converted JF669 and enables PALM imaging in the red emission channel. B) After JF669 is bleached, 405 nm light can be used to activate mEos2 and to excite the activated form with 561 nm light.

#### 6. PALM crosstalk imaging

HeLa cells were cultured in media containing Gibco fluorobrite Dulbecco's modified Eagle's medium (Thermo Fisher, cat. A1896701), Fetal Bovine Serum (Thermo Fisher, cat. 26140-079) 10%, sodium pyruvate (Thermo Fisher 11360-070) 1mM, Penicillin-Streptomycin (Thermo

Fisher, cat. 15140-122) 1%, and L-Glutamine (Thermo Fisher, cat. 25030-081) 4mM. 100 ng of each plasmid from Addgene (mEos2-Golgi-7 #57382 deposited by the Michael Davidson Lab) and (pcDNA5/FRT/TO_[Cox8a]x2_HaloTag7 #175529 deposited by the Kai Johnsson Lab) were used with GeneJet transfection reagent (SignaGen, cat. SL100488) to express mEos2-B4GALT on the Golgi and HaloTag on the inner mitochondrial membrane. After 24 h, cells were imaged in culturing media at 37 °C and 5% $CO_2$. EMCCD gain was set to 30, and a typical PALM shutter sequence was used, which consisted of 1 frame of 405 nm at 0.68 W/cm$^2$ light followed by 1 frame of 640 nm light at 680 W/cm$^2$ and 8 frames of 561 nm light at 480 W/cm$^2$. Localizations originating from pc-JF669 at the mitochondrial membrane were separated manually based on the overlap with the conventional far-red fluorescence signal, which is only detectable from the mitochondria and based on morphology. All other localizations are attributed to mEos2 originating from the Golgi based on the lack of co-localization with the conventional far-red fluorescence signal of JF669. In order to deplete oxygen at different rates, 0.2, 0.5, 1, and 2 µL of glucose oxidase from *Aspergillus niger* (Sigma-Aldrich, cat. G2133-10KU) and catalase from bovine liver (Sigma-Aldrich, cat.C9322-1G) stock solutions at 1000 U and 100 kU were added to samples before imaging.

#### 7. UV-free and additive-free PALM

To transiently transfect HeLa cells, 200 ng of pSEMS-TOM20-HaloTag plasmid (Addgene#111135) was used with GeneJet (SignaGen, cat. SL100488). After 24 h cells in 8 well chambered coverglass (Cellvis, cat. C8-1.5H-N) were incubated with 500 nM JF669-HaloTag ligand (Lavis Lab) to label the outer mitochondrial membrane. After washing three times with PBS, cells were incubated in 300µL of growth media before cells were imaged with alternating 640 nm activation light and 561 nm excitation light at 37 °C and 5% $CO_2$. After 5000 frames, single-molecule localizations were rendered as a 2D Gaussian whose width is weighted by the inverse square root of the detected number of photons. For single-molecule tracking analysis, the single-molecule localizations that appeared within 0.5 µm in more than 5 consecutive 561 nm frames (50 ms exposure) were linked to a trace.

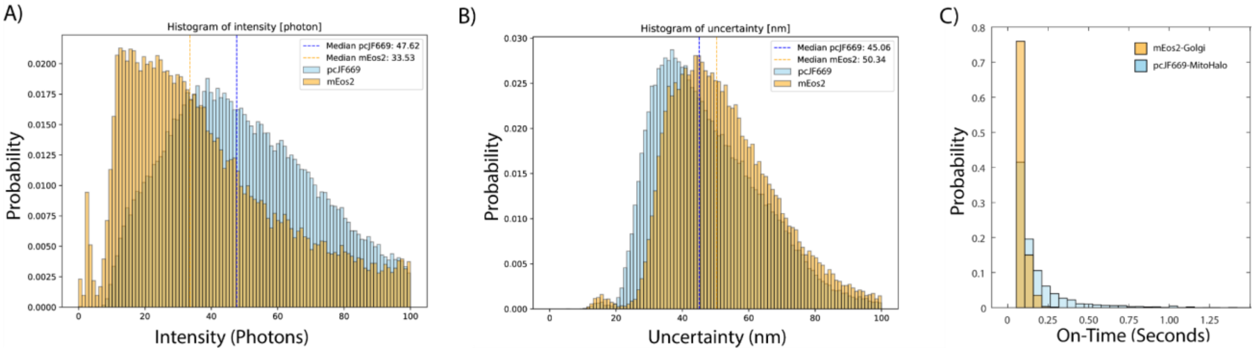


Figure S8: A) Probability distribution of photons per localization of different labels mEos2-Golgi (orange) and pc-JF669 HaloTag-TOM20 (blue). B) Probability distribution of uncertainty of different labels mEos2-Golgi (orange) and pc-JF669 HaloTag-TOM20 (blue) C) Probability distribution of on-times different labels mEos2-Golgi (orange) and pc-JF669 HaloTag-TOM20 (blue). pc-JF669 is brighter and more photo-stable than mEos2, leading to higher resolution and longer traces.

#### 8. Emission channel transformation

To precisely overlay the two microscope channels in two-color imaging experiments, the transformation between the two has to be determined as described in a previous publication[23]. In short, fluorescein at 10µM in DI water was mounted on a coverslip and a 9x9 grid of diffraction-limited points was projected onto the sample plane with 405 nm illumination using a digital mirror device. Grid points were localized with insight and a 3rd order polynomial transformation was generated to map top channel localizations to the bottom channel. This error had an average error of 16 nm, on par with localization uncertainty.

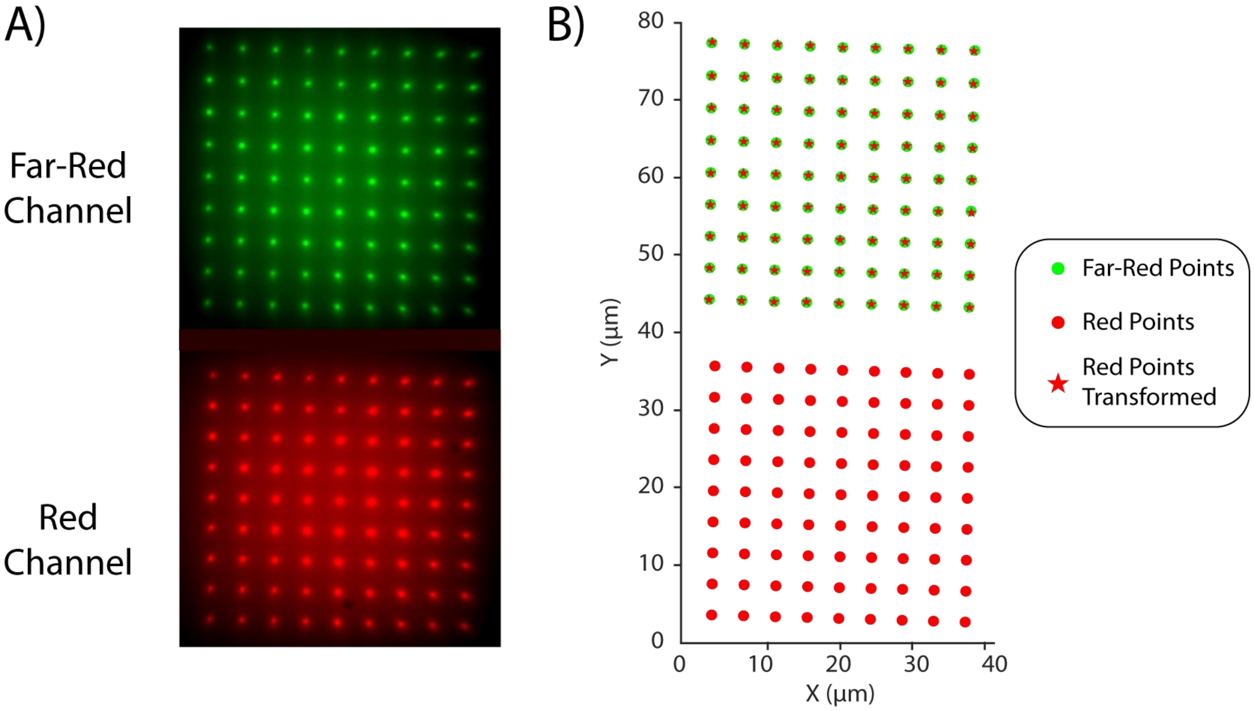


Figure S9. A) A grid of points was projected onto fluorescein on the sample plane with 405 nm excitation to produce a grid of emissions in both emission channels. B) Those points were then localized and a 3rd order polynomial transformation was generated to transform localization from the red channel to the far-red channel with an average error of 16 nm, which is approximately the localization precision.

#### 9. Pseudo two-Color PALM in the same emission channel

To transiently transfect HeLa cells, 100 ng of each, the Tom20-HaloTag plasmid and the mEos2-B4GALT1 plasmid were used with GeneJet. After 24 h, cells were incubated with 500 nM JF669-HaloTag ligand to label the mitochondria. Cells were then washed 3 times and imaged at 37 °C and 5% $CO_2$ in media containing Gibco fluorobrite Dulbecco's modified Eagle's medium (Thermo Fisher, cat. A1896701), Fetal Bovine Serum (Thermo Fisher, cat. 26140-079) 10%, sodium pyruvate (Thermo Fisher 11360-070) 1mM, Penicillin-Streptomycin (Thermo Fisher, cat. 15140-122) 1%, and L-Glutamine (Thermo Fisher, cat. 25030-081) 4mM. Cells were first imaged with alternating 640 nm activation light and 561 nm excitation light to image and bleach all of the JF669 labeling the outer mitochondrial membrane. After all JF669 was bleached, alternating 405 nm activation light and 561 nm excitation light was used to image mEos2 on the Golgi. Localizations from the red emission channel were rendered as a 2D Gaussian whose width is weighted by the inverse square root of the detected number of photons.

#### 10. TXTL

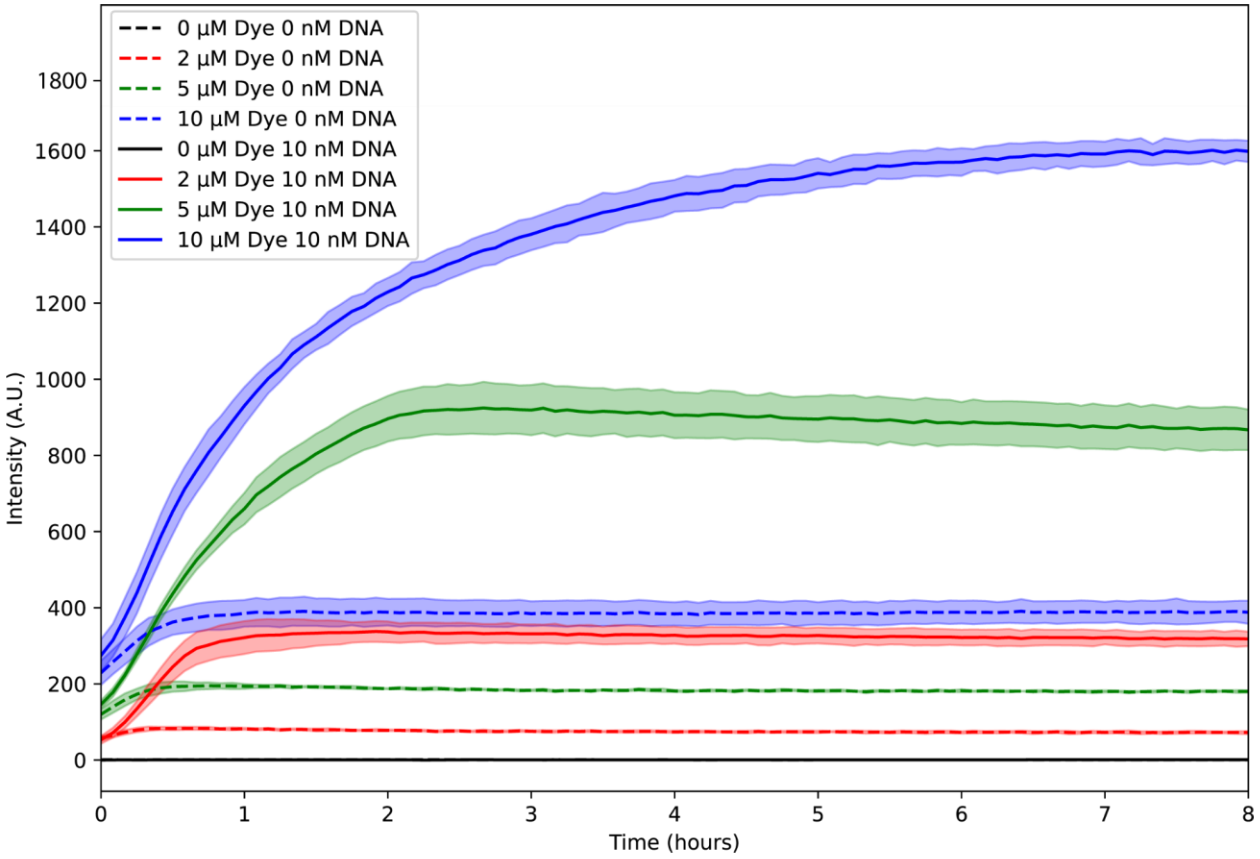


Figure S10. 620 nm/670 nm E kinetics of a blank TXTL with no DNA (dashed line), and 10 nM of DNA coding for our Halotag construct, p70a-HaloTag-MSA (solid line). The reactions were incubated with 0 µM (black), 2 µM (red), 5 µM (green), and 10 µM (blue)

of the JFX650 dye. Samples without the DNA showed relatively constant fluorescence, but samples with the HaloTag construct show an increase in fluorescence due to the fluorogenic dye binding to the HaloTag protein. With 5 µM dye, the fluorescence saturates at around 2 h, showing that the overwhelming majority of dyes are bound to HaloTag at this time. The concentration of HaloTag is therefore approximately 5 µM after 2 h in this experiment.

**Cell-free transcription-translation.** Cell-free gene expression was carried out using an *E. coli* TXTL system described previously[36], with one modification. We used the strain BL21-Δ*recBCD* Rosetta2 in which the *recBCD* gene set is knocked out to prevent the degradation of linear DNA[37]. The preparation and usage of the TXTL system were the same as reported before[36]. Briefly, *E. coli* cells were grown in a 2xYT medium supplemented with phosphates. Cells were pelleted, washed, and lysed with a cell press. After centrifugation, the supernatant was recovered and pre-incubated at 37 °C for 80 min. After a second centrifugation step, the supernatant was dialyzed for 3 h at 4 °C. After a final spin-down, the supernatant was aliquoted and stored at −80 °C. The TXTL reactions comprised the cell lysate, the energy and amino acid mixtures, maltodextrin (30 mM) and ribose (30 mM), magnesium (2-5 mM) and potassium (50-100 mM), PEG8000 (1-2 wt%), water and the DNA to be expressed. The TXTL reactions were incubated at 30 °C on 96 well plates (2 µl reactions) for the measurement of the kinetics of HaloTag synthesis.

#### 11. *In vitro* sample preparation, imaging and analysis

Fisherbrand #1 25x25-1 coverslips (Thermo Fisher, cat. 12-542C) were first washed with 2% Micro 90 (International Products Corporation, cat. 89210-140-EA) before being rinsed with DI water and dried with compressed nitrogen. Dried coverslips were plasma cleaned in a plasma cleaner (Harrick, cat. PDC-32G) on the low setting for 10 minutes. Plasma-cleaned coverslips were submerged in Biotinylated Bovine Serum Albumin (Biotin-LC-BSA) (3 biotin/BSA) (Abcam, cat. ab286862) for 1 h. TXTL was incubated for 2 h with 10 nM of linear P70a-MSA-HaloTag DNA (Twist, sequence:
CACCATCAGCCAGAAAACCGAATTTTGCTGGGTGGGCTTCCGCTGGGCATGCTGAGCTAA
CACCGTGCGTGTTGACAATTTTACCTCTGGCGGTGATAATGGTTGCAGCTAGCAATAATTTT
GTTTAACTTTAAGAAGGAGATATACCATGGCCGAAGCCGGTATCACCGGCACCTGGTACAA
CCAGTCTGGTTCTACCTTCACCGTTACCGCGGGTGCGGACGGTAACCTGACCGGTCAGTA
CGAAAACCGTGCGCAGGGCACTGGTTGCCAGAACTCTCCGTACACCCTGACCGGTCGTTA
CAACGGTACCAAACTGGAATGGCGTGTTGAATGGAACAACTCTACCGAAAACTGCCACTCT
CGTACCGAATGGCGTGGTCAGTACCAGGGTGGTGCGGAAGCGCGTATCAACACCCAGTG
GAACCTGACCTACGAAGGTGGTTCTGGTCCGGCGACCGAACAGGGTCAGGACACCTTCAC
CAAAGTTAAAGGGCCCTCCGGACTCAGATCTCGAGCTgcagaaatcggtactggctttccattcgaccccca
ttatgtggaagtcctgggcgagcgcatgcactacgtcgatgttggtccgcgcgatggcacccctgtgctgttcctgcacggtaacccga
cctcctcctacgtgtggcgcaacatcatcccgcatgttgcaccgacccatcgctgcattgctccagacctgatcggtatgggcaaatcc
gacaaaccagacctgggttatttcttcgacgaccacgtccgcttcatggatgccttcatcgaagccctgggtctggaagaggtcgtcctg
gtcattcacgactggggctccgctctgggtttccactgggccaagcgcaatccagagcgcgtcaaaggtattgcatttatggagttcatc
cgccctatcccgacctgggacgaatggccagaatttgcccgcgagaccttccaggccttccgcaccaccgacgtcggccgcaagct
gatcatcgatcagaacgtttttatcgagggtacgctgccgatgggtgtcgtccgcccgctgactgaagtcgagatggaccattaccgcg

agccgttcctgaatcctgttgaccgcgagccactgtggcgcttcccaaacgagctgccaatcgccggtgagccagcgaacatcgtcgcgctggtcgaagaatacatggactggctgcaccagtcccctgtcccgaagctgctgttctggggcaccccaggcgttctgatcccaccggccgaagccgctcgcctggccaaaagcctgcctaactgcaaggctgtggacatcggcccgggtctgaatctgctgcaagaagacaacccggacctgatcggcagcgagatcgcgcgctggctgtcgacgctggagatttccggctaaCTCGAGCAAAGCCCGCCGAAAGGCGGGCTTTTCTGTGTCGACCGATGCCCCGCTTCCTCGCTCACTG), before being diluted 500x in PBS (Thermo Fisher, cat. 10010023) with JFX650-HaloTag Ligand (Lavis Lab) at a concentration of 10 nM. Coverslips were rinsed with PBS before 10 µL of this 10 nM solution of dye plus HaloTag construct was placed on top of the biotinylated surface. After 1 h, coverslips were rinsed three times with PBS to wash away TXTL, unbound dyes, and unbound proteins. To seal *in vitro* samples, Scientific Gene Frame stickers (Thermo Fisher, cat. AB0577) were placed on cleaned glass slides and coverslips with 10 µL of PBS were then sealed for imaging. Imaging was performed for 5000 frames at 20hz with EMCCD gain 30. The laser shutter sequence was composed of 19 frames of 640 nm illumination at 6.8 kW/cm$^2$ followed by a single frame of 561 nm illumination at 6.8 kW/cm$^2$ . This sequence was repeated for the duration of the movie. SMLM analysis was performed using the INSIGHT software (Zhuang lab, Harvard), and cross-validated using the ThunderSTORM plugin for ImageJ (Fiji). Single-molecule recognition was confirmed by visual perception of fluorescent blinking, and single-molecule identification parameters for 2D Gaussian PSFs were set accordingly (Gaussian height ≥ 50 photons, width (200–750) nm, ROI: 7 × 7 pixels). All super-resolution images were represented across at least 5000 frames.

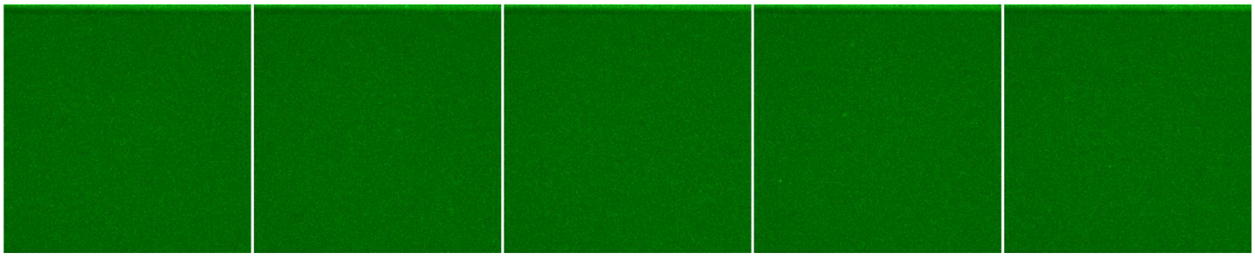

Figure S11. Plasma-cleaned coverslips covered with Biotin-BSA and TXTL solution with no dye were washed 3 times with PBS before imaging. Five random frames from five random samples are displayed to show minimal single-molecule autofluorescence in the red channel.
